# Pax6-dependent patterning in an annelid informs the evolution of bilaterian nerve cords

**DOI:** 10.64898/2026.06.27.734823

**Authors:** Jovana Doderovic, Miroslava Kolek, Anna Zitova, Iryna Kozmikova, Zbynek Kozmik

## Abstract

Conserved dorsoventral patterning systems have been proposed as evidence for a common evolutionary origin of centralized nervous systems in Bilateria, yet functional evidence outside vertebrates and arthropods remains limited. Here, we investigated the role of *pax6* in the annelid *Platynereis dumerilii* using a mutant carrying a 61 bp deletion in the paired-domain coding region. Loss of pax6 disrupted ventral neuroectodermal patterning at 34 hpf, causing a shift in *nk2.2* expression, narrowing of the *nk6* domain, and downregulation of *pax3/7*, while *msx* expression remained largely unaffected. These early patterning defects were followed by selective neuronal abnormalities at 48 hpf, including displacement of *TrpH*-positive serotonergic cells and loss of posterior *hb9*-positive motoneuron domains. By 6 dpf, additional defects were observed in *TrpH*, *ChAT*, *VAChT*, and *nk2.2* expression, accompanied by severe disruption of ventral nerve cord morphology and loss of the characteristic rope-ladder architecture. Together, these findings identify pax6 as a key regulator linking dorsoventral progenitor patterning, neuronal subtype specification, and nervous system morphogenesis in *Platynereis*. Our results provide functional evidence that the conserved dorsoventral patterning network plays an essential role in annelid ventral nerve cord development and support the view that important components of bilaterian nervous system patterning predate the divergence of major animal lineages.

## 1. Introduction

Nervous systems exhibit remarkable diversity across Metazoa, ranging from diffuse nerve nets in cnidarians to highly centralized systems composed of brains and longitudinal nerve cords in bilaterians (Adameyko, 2023; Arendt et al., 2008; Arendt et al., 2016; Martín-Durán and Hejnol, 2021; Watanabe et al., 2009). The evolutionary origin of centralized nervous systems remains one of the central questions in evolutionary developmental biology. Although some groups, such as Xenacoelomorpha, possess comparatively simple or weakly centralized nervous systems, centralization is widespread throughout Bilateria, raising the possibility that at least some aspects of nervous system organization were already present in the bilaterian ancestor (Arendt et al., 2016; Brunet and Arendt, 2016; Cannon et al., 2016).

One influential hypothesis proposes that centralized nervous systems in bilaterians share a common evolutionary origin. Support for this idea comes from the remarkable conservation of developmental mechanisms underlying nervous system patterning. In vertebrates, arthropods, and annelids, neural tissue develops opposite the major source of BMP signaling, and the neuroectoderm is regionalized by homologous sets of transcription factors expressed in staggered dorsoventral domains (Arendt and Nübler-Jung, 1999; De Robertis and Kuroda, 2004; Denes et al., 2007). These domains are defined by members of the Nkx, Pax, and Msx transcription factor families and give rise to distinct neuronal populations along the mediolateral axis of the developing nervous system (Briscoe and Novitch, 2008; Briscoe et al., 2000; Jessell, 2000; Pattyn et al., 2003). In vertebrates and insects, extensive cross-regulatory interactions among neighboring domains establish sharp expression boundaries and contribute directly to neuronal subtype specification (Crews, 2019; Ericson et al., 1997; Sander et al., 2000).

However, the interpretation of these similarities remains controversial. Comparative studies in xenacoelomorphs, brachiopods, rotifers, and several spiralian taxa have revealed cases in which dorsoventral patterning genes are expressed without obvious correspondence to differentiated neuronal populations (Beckers et al., 2019; Carrillo-Baltodano et al., 2021; Martín-Durán et al., 2018). Such observations have been interpreted as evidence that nerve cords may have evolved independently in different bilaterian lineages and that similarities in gene expression patterns could reflect repeated co-option of ancient developmental programs rather than strict homology (Arendt, 2018; Martín-Durán et al., 2018). Consequently, expression data alone is insufficient to resolve the debate. Functional studies testing whether these genes perform comparable developmental roles across distantly related animals are needed.

The marine annelid *Platynereis dumerilii* provides an attractive system for such analyses. *Platynereis* possesses a centralized nervous system consisting of a brain and a rope-ladder ventral nerve cord and is widely regarded as retaining many ancestral bilaterian characteristics (Arendt et al., 2008; Denes et al., 2007; Özpolat et al., 2021). Previous studies demonstrated that the ventral neuroectoderm of *Platynereis* is patterned by staggered mediolateral expression domains of *nk2.2*, *nk6*, *pax6*, *pax3/7*, and *msx*, closely resembling the arrangement observed in vertebrates and insects (Denes et al., 2007). Moreover, these domains correlate with distinct neuronal populations, including serotonergic, cholinergic, sensory, and interneuron classes. Nevertheless, whether this conserved molecular organization reflects a shared functional mechanism has remained unknown.

Among the components of this network, *pax6* is particularly well suited for functional investigation. Pax6 is an evolutionarily conserved paired-box transcription factor that plays fundamental roles in the development of eyes, the central nervous system, endocrine organs, and sensory structures throughout Bilateria (Callaerts et al., 1997; Hill et al., 1991; Walther and Gruss, 1991). In vertebrates, *Pax6* occupies an intermediate position within the dorsoventral patterning network of the neural tube and participates in reciprocal interactions with Nkx2.2 and Nkx6-class transcription factors that establish progenitor-domain boundaries and regulate neuronal subtype specification (Briscoe et al., 2000; Ericson et al., 1997; Vallstedt et al., 2001). Similar functions have been described in insects, where *Pax6* homologs contribute to nervous system development in addition to their well-known role in eye formation (Callaerts et al., 2001; Furukubo-Ttokunaga et al., 2009). Whether *Pax6* performs comparable patterning functions in annelids remains largely unexplored.

In this study, we generated a stable *pax6* knockout line in *Platynereis dumerilii* using zinc-finger nuclease-mediated genome editing and examined the consequences of *pax6* loss on ventral nerve cord development. We analyzed the expression of dorsoventral patterning genes (*nk2.2*, *nk6*, *pax3/7*, and *msx*), markers of differentiated neuronal populations (*TrpH*, *Hb9*, *ChAT*, and *VAChT*), and overall nervous system morphology. Our results demonstrate that *pax6* is required for proper organization of dorsoventral progenitor domains, neuronal subtype specification, and maturation of ventral nerve cord architecture. These findings provide functional evidence that the conserved dorsoventral patterning system identified in annelids is developmentally significant and contributes to broader discussions concerning the evolutionary origins of centralized nervous systems in Bilateria.

## 2. Materials and Methods

### 2.1. Animal culture and whole mount in situ hybridization

*Platynereis dumerilii* breeding cultures were maintained as previously described (Kuehn et al., 2019). Embryos at selected developmental stages were fixed for 2 h at room temperature in 4% paraformaldehyde and stored in methanol at -20°C. Whole-mount in situ hybridization was performed as described previously (Žídek et al., 2018). Embryos were permeabilized with Proteinase K (Roche) for 1 min at 34 and 48 hpf and for 2.5 min at 6 dpf. Hybridization with digoxigenin-UTP-labeled RNA probes was performed at 63°C. Samples were then incubated with anti-digoxigenin alkaline phosphatase-conjugated antibody (Roche), and signal was developed using the Vector Blue Alkaline Phosphatase Substrate Kit (Vector Laboratories). Acetylated tubulin was detected using a mouse monoclonal anti-acetylated tubulin antibody (1:500; Sigma, T6793), followed by incubation with a fluorescent secondary antibody (Invitrogen, 1:500). Samples were mounted in TDE and imaged using a Leica TCS SP8 confocal microscope. Images were processed in Fiji.

### 2.2. Generation and maintenance of Pax6 knockout line

To design pairs of Zinc Finger Nucleases (ZFNs) the paired domain exon *Platynereis dumerilii pax6* (AM114770.1) was selected. Three pairs of ZFNs were custom designed and generated (Sigma-Aldrich). ZFN pair recognizing binding site TGCCCCTCCATTTTC<u>gcctg</u>GGAAATAAGGGACCG was chosen for genome editing (cut site underlined). ZFN mRNA in a concentration of 50 ng/μl was injected into fertilized eggs of *Platynereis* wild-type parents. The eggs were kept at 18°C for 45 min before injection and were injected at 14.5°C. The injected individuals were kept at 18°C for 5 to 8 days in 6-well-plates (Nunc multidish no. 150239, Thermo Scientific) and then cultured at 22°C until sexual maturity. The mature worms were crossed to wild-type worms and the progeny was genotyped, resulting in one founder line, which carried 61bp frameshift-causing deletion. Animals were genotyped using primers TTTCAGGTGTCGAACGGTTGT and ACAACAGCCTGTCCCCATATC. Due to lethality of homozygous pax6 mutation, the line was maintained as heterozygotes and all experiments were performed on progenies of heterozygote crosses.

## 3. Results

### 3.1. Generation and validation of a pax6 knockout line

To investigate the role of pax6 during ventral nerve cord development, we generated a loss-of-function mutant line in *Platynereis dumerilii* using zinc-finger nuclease (ZFN)-mediated genome editing. A ZFN pair was designed to target the coding sequence of the paired domain of pax6 (Fig. 1A). Sequence analysis identified a mutant allele carrying a 61 bp deletion at the target site (Fig. 1B). This deletion introduces a premature stop codon and is predicted to generate a severely truncated pax6 protein lacking essential functional domains.

**Fig. 1.**
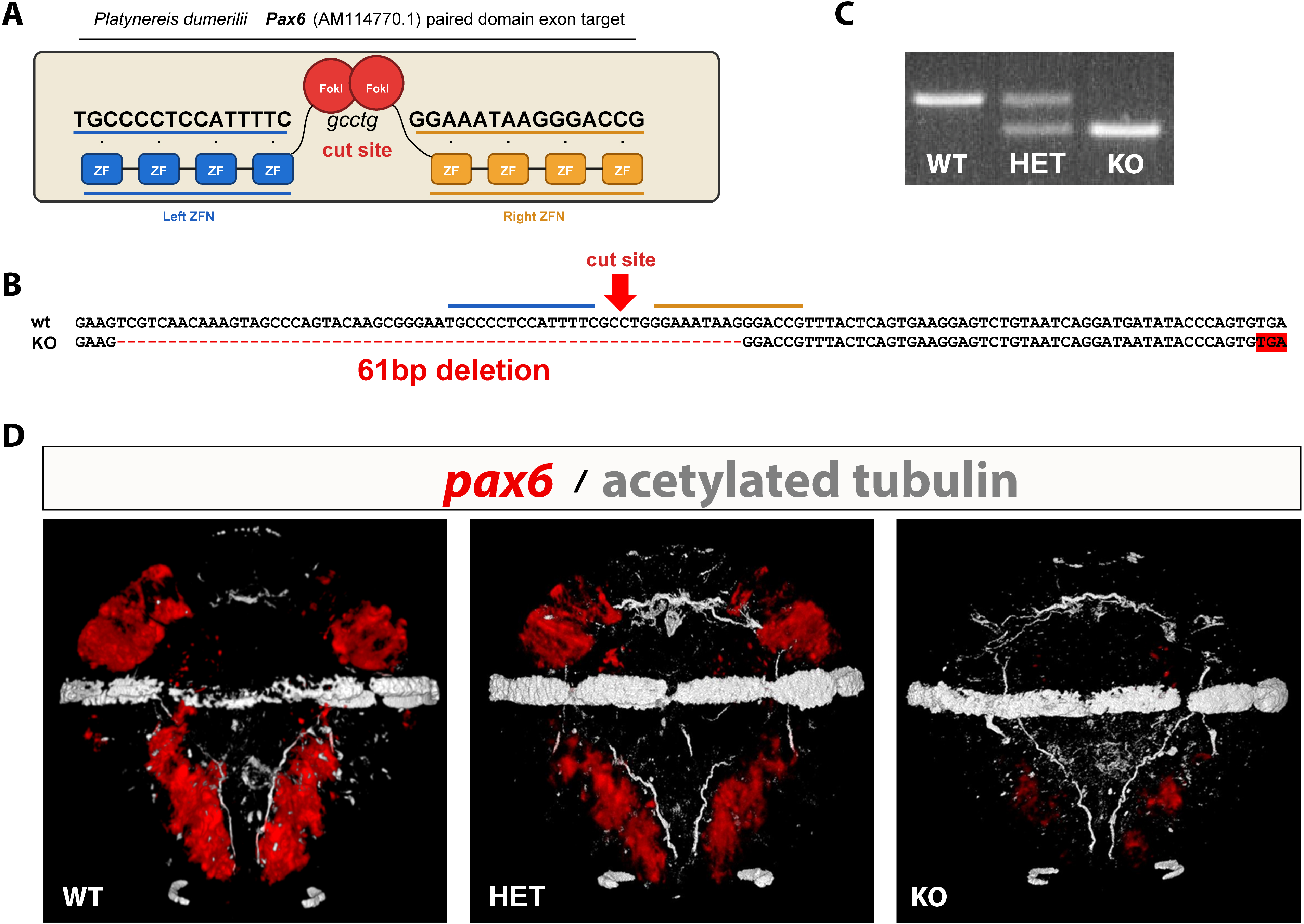
Generation and validation of a *pax6* knockout line in *Platynereis dumerilii*. (A) Schematic representation of the zinc-finger nuclease (ZFN) strategy used to target the paired-domain coding exon of *pax6* (AM114770.1). Left and right ZFNs bind adjacent target sequences and direct FokI-mediated cleavage at the indicated cut site. (B) Sequence analysis of the mutant allele showing a 61 bp deletion at the ZFN target site. The deletion introduces a premature stop codon (red box), predicted to truncate the pax6 protein and trigger nonsense-mediated decay of the mutant transcript. (C) Genotyping PCR of WT, heterozygous (HET), and homozygous knockout (KO) animals. The KO allele is distinguished by the reduced amplicon size resulting from the 61 bp deletion, while HET animals display both WT and mutant bands. (D) Whole-mount in situ hybridization (WMISH) for *pax6* in WT, HET, and KO larvae. WMISH signal is shown in red and acetylated tubulin immunostaining is shown in white. *pax6* expression is absent in KO animals. Representative ventral views of 34 hpf larvae are shown.

Genotyping PCR readily distinguished wild-type (WT), heterozygous (HET), and homozygous knockout (KO) animals based on the amplicon-size difference caused by the deletion (Fig. 1C). To assess the effect of the mutation on pax6 expression, we performed whole-mount in situ hybridization (WMISH). Strong pax6 expression was detected in WT and HET larvae, whereas the signal was undetectable in KO animals (Fig. 1D), confirming successful disruption of the *pax6* locus. These results establish a stable *pax6* knockout line suitable for functional analysis of ventral nerve cord patterning and neuronal differentiation.

### 3.2. Pax6 loss alters mediolateral patterning domains at 34 hpf

At 34 hpf, we analyzed the expression patterns of the dorsoventral patterning genes *nk2.2*, *nk6*, *pax3/7* and *msx* (Fig. 2). In WT embryos, *nk2.2* expression was detected in the medial trunk region and followed the ventral nerve cord connectives (Fig. 2A). In pax6KO embryos, the anterior *nk2.2*-positive territory was shrinked in the posterior region and the boundaries of the expression domain were altered (Fig. 2B).

**Fig. 2.**
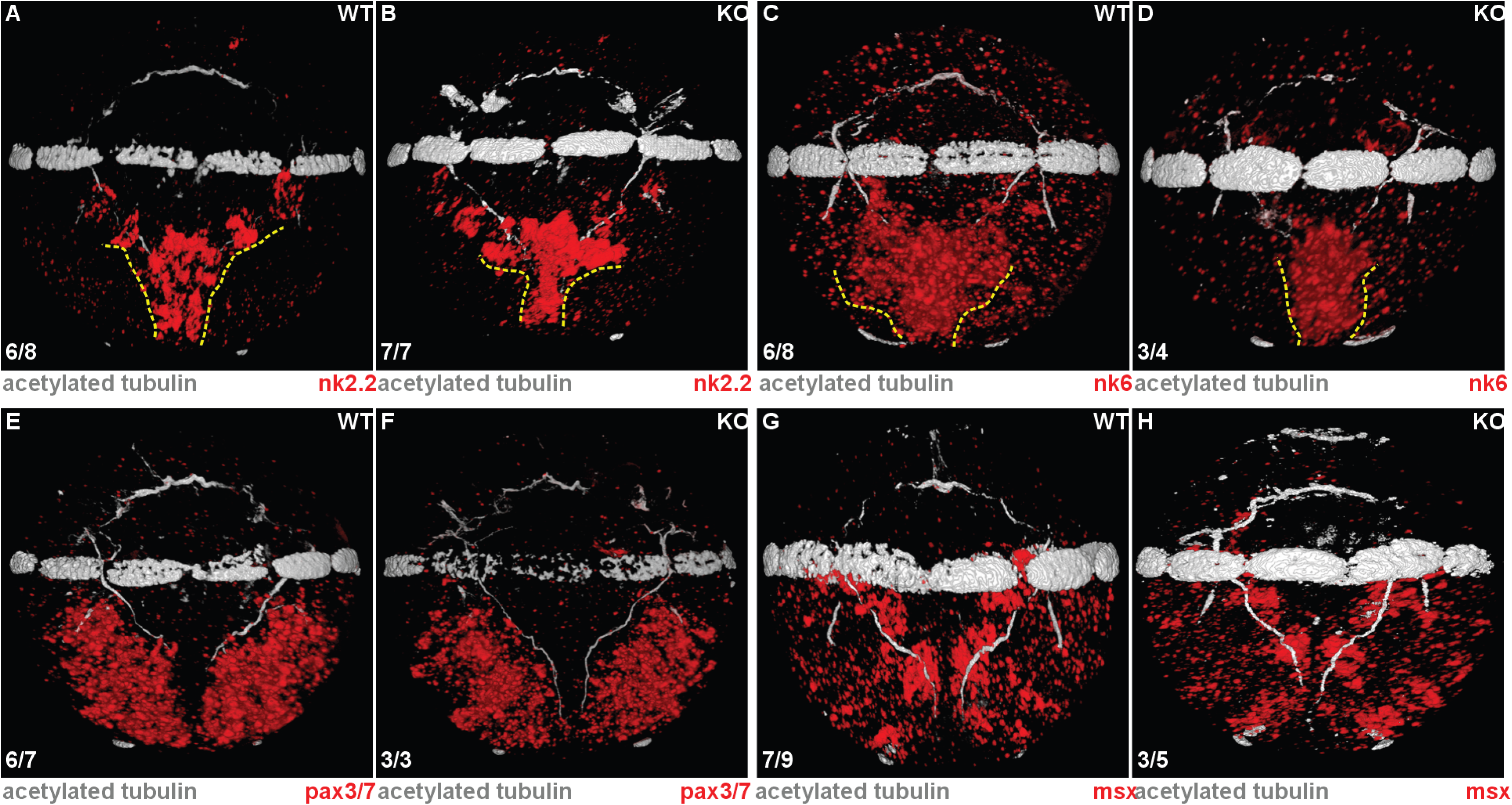
Whole-mount in situ hybridization (WMISH) analysis of ventral nerve cord patterning at 34 hpf. Expression of *nk2.2* (A,B), *nk6* (C,D), *pax3/7* (E,F), and *msx* (G,H) in WT (A,C,E,G) and KO (B,D,F,H) embryos. WMISH signal is shown in red. The ventral nerve cord (VNC) is visualized by acetylated tubulin immunostaining (white). Yellow dashed lines in A–D outline altered boundaries of the *nk2.2* and *nk6* expression domains in KO embryos relative to WT. Ventral substacks are shown; all images are ventral views. Numbers in the lower left corner indicate the number of embryos displaying the representative phenotype out of the total number analyzed.

The *nk6* domain was also affected by loss of *pax6*. In WT embryos, *nk6* was expressed broadly across the ventral trunk region (Fig. 2C), whereas in pax6KO embryos this domain was constricted into a narrower medial stripe associated with the ventral nerve cord connectives (Fig. 2D). Expression of *pax3/7* was reduced in pax6KO embryos, although the overall lateral position of the domain was retained (Fig. 2E,F). By contrast, no consistent change in *msx* expression was detected in pax6KO embryos (Fig. 2G,H). Together, these data indicate that *pax6* is required for proper mediolateral organization of the ventral neuroectoderm and preferentially affects domains adjacent to, or overlapping with, the *pax6* expression territory.

### 3.3. Early patterning defects are followed by selective neuronal marker abnormalities at 48 hpf

To determine whether the early patterning defects were associated with altered neuronal differentiation, we examined markers of specific neuronal populations at 48 hpf, including *TrpH*, *ChAT*, *hb9* and *VAChT* (Fig. 3). The serotonergic neuron marker *TrpH* showed a reproducible change in pax6KO embryos. In WT embryos, two posterior *TrpH*-positive domains were positioned bilaterally above the ventral nerve cord connectives near the level of the third commissure (Fig. 3A). In pax6KO embryos, these domains were displaced toward the midline of the ventral nerve cord (Fig. 3B).

**Fig. 3.**
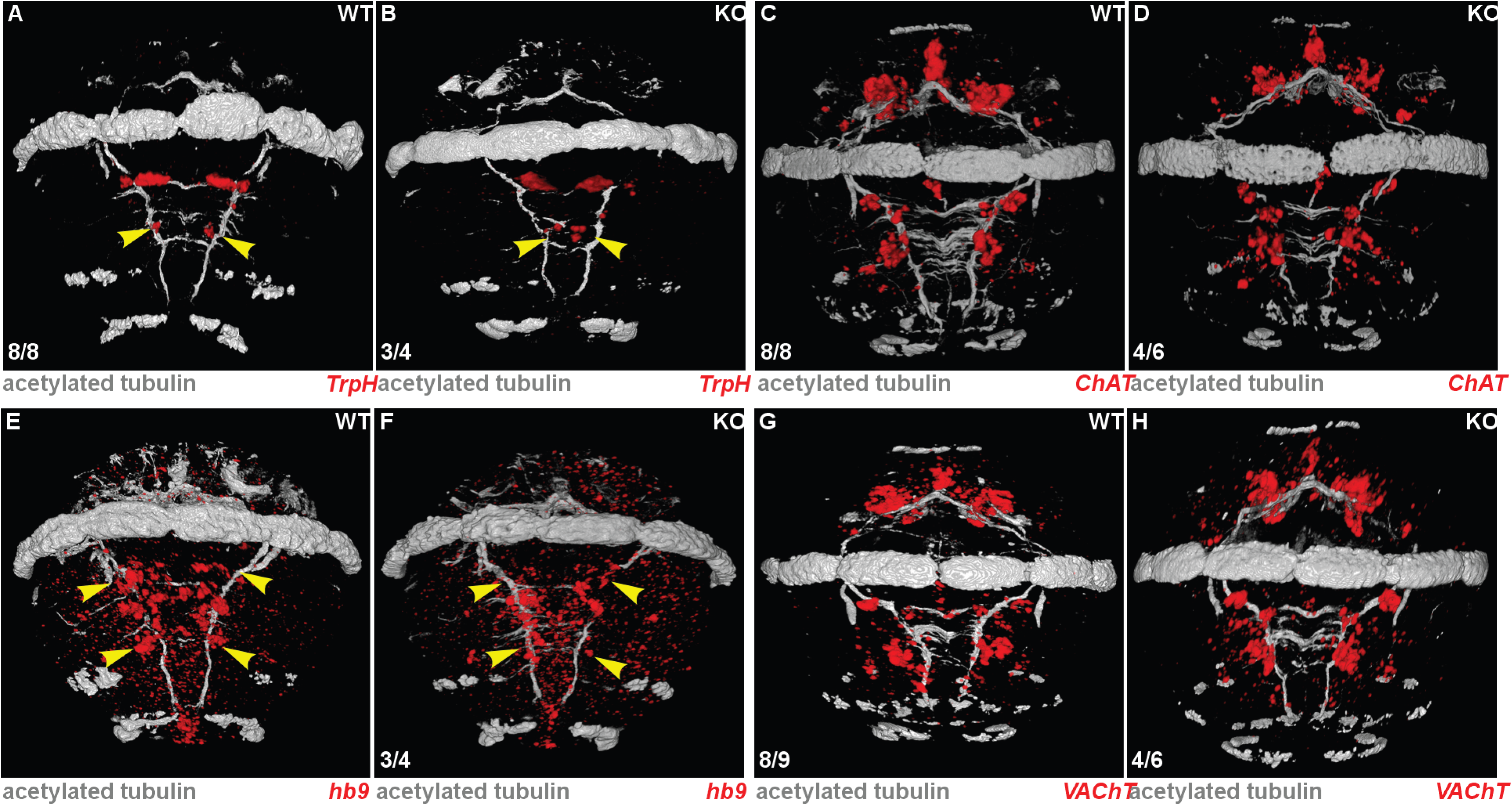
Whole-mount in situ hybridization (WMISH) analysis of neuronal marker expression at 48 hpf. Expression of *TrpH* (A,B), *ChAT* (C,D), *hb9* (E,F), and *VAChT* (G,H) in WT (A,C,E,G) and KO (B,D,F,H) embryos. WMISH signal is shown in red. The ventral nerve cord (VNC) is visualized by acetylated tubulin immunostaining (white). Yellow arrowheads in A,B and E,F indicate altered *TrpH* and *hb9* expression domains in KO embryos relative to WT. Ventral substacks are shown; all images are ventral views. Numbers in the lower left corner indicate the number of embryos displaying the representative phenotype out of the total number analyzed.

The motoneuron-associated marker *hb9* was also affected. In WT embryos, posterior *hb9*-positive domains were detected along the ventral nerve cord (Fig. 3E). These posterior domains were absent or strongly reduced in pax6KO embryos (Fig. 3F). In contrast, no obvious changes were detected in *ChAT* or *VAChT* expression at this stage (Fig. 3C,D,G,H). These results suggest that loss of pax6 does not cause a uniform delay in neuronal differentiation, but instead selectively affects neuronal populations associated with altered dorsoventral progenitor territories.

### 3.4. Neuronal marker defects become more pronounced at 6 dpf

At 6 dpf, when the larval nervous system is highly differentiated, we examined the expression of *TrpH*, *ChAT*, *VAChT* and *nk2.2* (Fig. 4). In pax6KO larvae, *TrpH*-positive cells along the ventral nerve cord were reduced or absent, particularly near the second and third commissural regions (Fig. 4G,H). *ChAT* and *VAChT* expression domains were also altered in pax6KO larvae, appearing broader and less precisely organized than in WT controls (Fig. 4A-D). The nk2.2 expression pattern remained abnormal at 6 dpf. In pax6KO larvae, *nk2.2*-positive domains near the first and second commissures were enlarged and displaced, and the domain near the third commissure was altered (Fig. 4E,F). Expression patterns of *nk6*, *pax3/7*, *msx* and *hb9* were not examined at this stage. Together, the 6 dpf data indicate that the early patterning defects observed at 34 hpf persist and are accompanied by later abnormalities in neuronal marker organization.

**Fig. 4.**
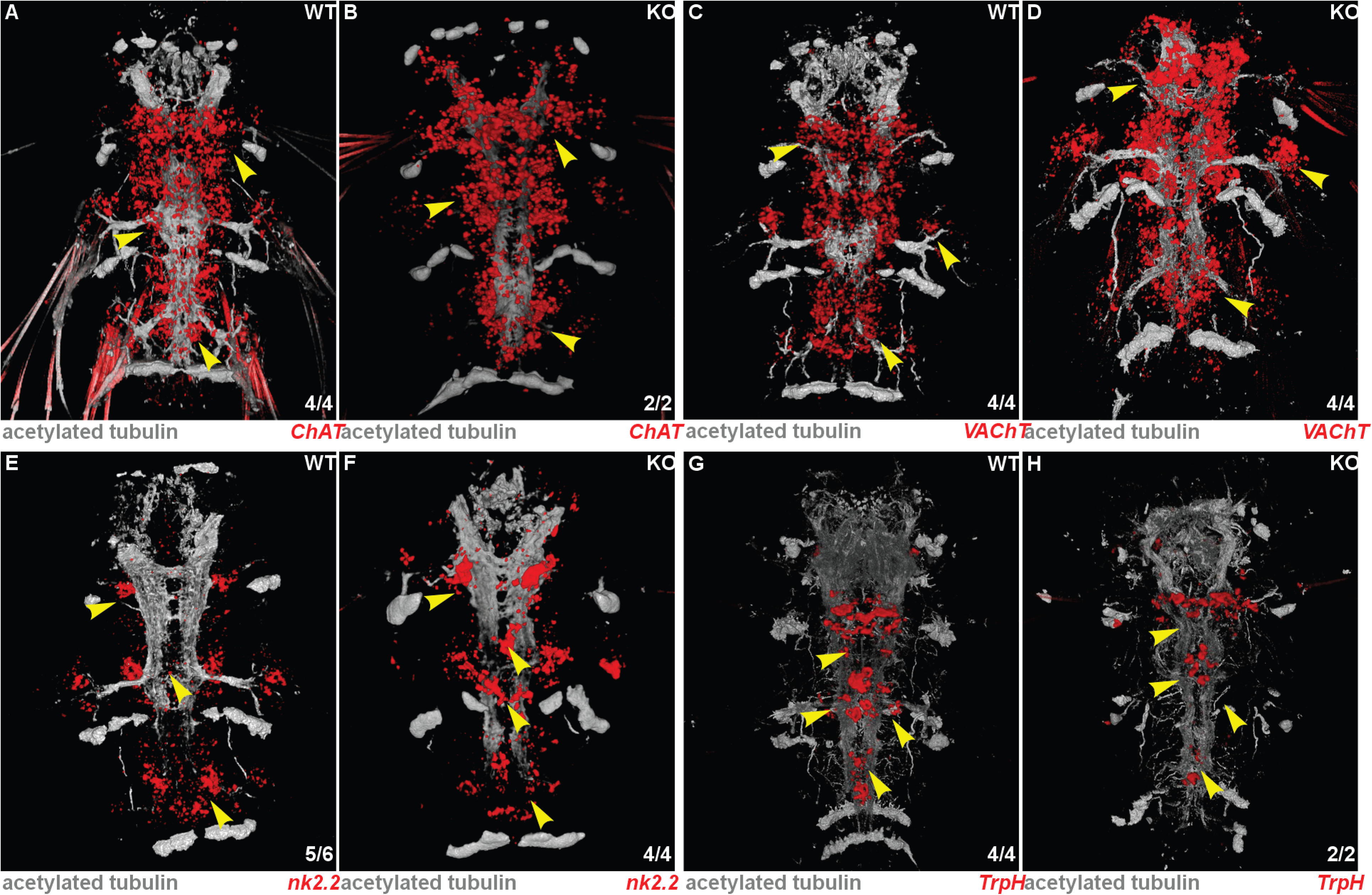
Whole-mount in situ hybridization (WMISH) analysis of neuronal and progenitor marker expression at 6 dpf. Expression of *ChAT* (A,B), *VAChT* (C,D), *nk2.2* (E,F), and *TrpH* (G,H) in WT (A,C,E,G) and KO (B,D,F,H) larvae. WMISH signal is shown in red. The ventral nerve cord (VNC) is visualized by acetylated tubulin immunostaining (white). Yellow arrowheads highlight regions showing altered *ChAT, VAChT, nk2.2*, and *TrpH* expression in KO larvae relative to WT controls. Ventral substacks are shown; all images are ventral views. Numbers in the lower right corner indicate the number of larvae displaying the representative phenotype out of the total number analyzed.

### 3.5. Loss of *pax6* disrupts ventral nerve cord morphology during larval development

To determine whether the molecular defects were accompanied by broader abnormalities in nervous system organization, we examined acetylated tubulin staining at successive developmental stages (Fig. 5). At 34 hpf and 48 hpf, the overall architecture of the ventral nerve cord appeared largely preserved in pax6KO animals, despite the changes in dorsoventral patterning and neuronal marker expression described above (Fig. 5A-D). In contrast, pronounced morphological abnormalities became evident by 6 dpf (Fig. 5E,F).

**Fig. 5.**
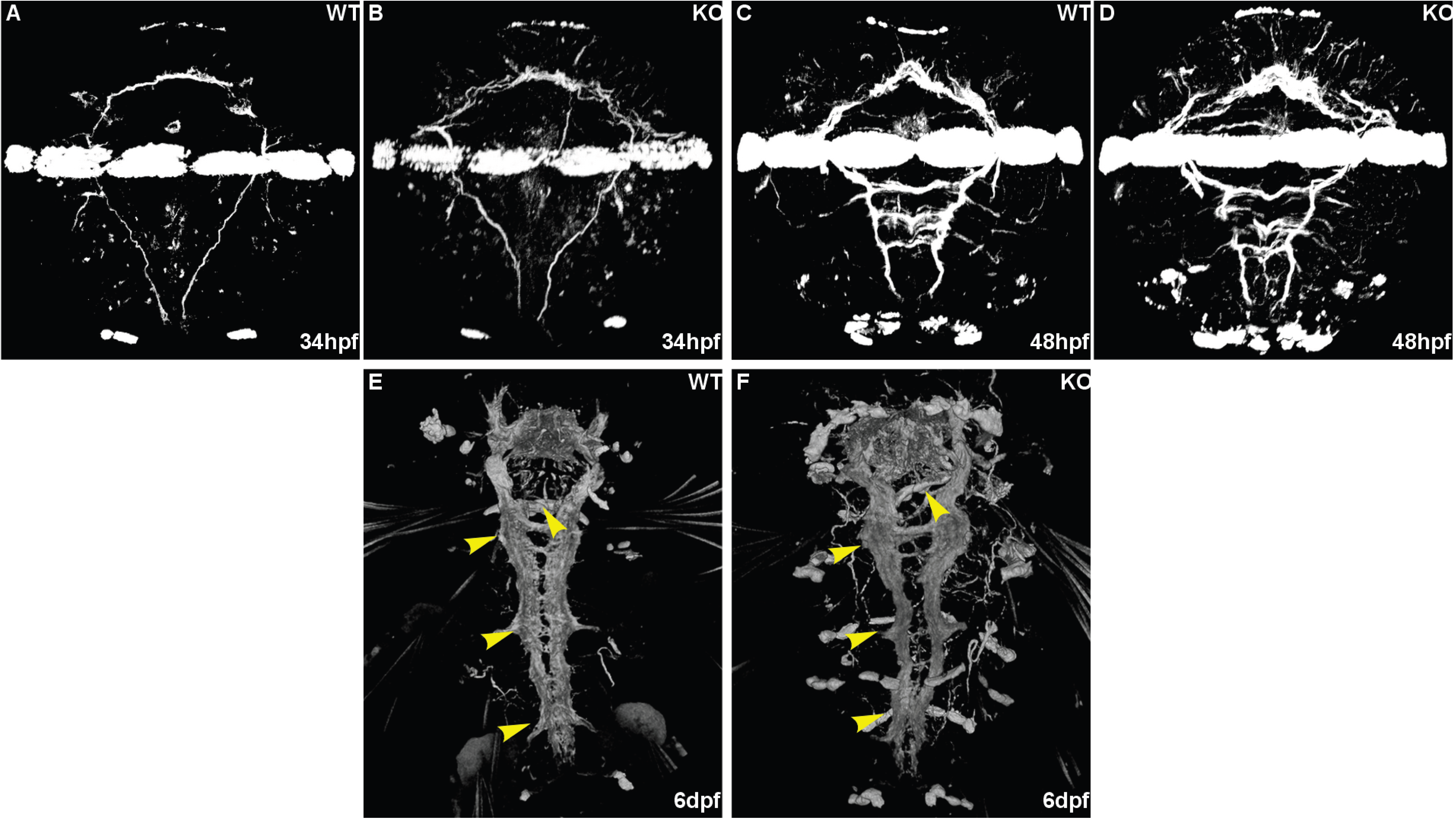
Acetylated tubulin immunostaining reveals overall nervous system morphology in WT and *pax6* KO animals. Representative ventral views of WT (A,C,E) and KO (B,D,F) animals at 34 hpf (A,B), 48 hpf (C,D), and 6 dpf (E,F). Acetylated tubulin signal is shown in white. Yellow arrowheads in E and F highlight morphological differences in the ventral nerve cord of KO animals relative to WT controls. All images are ventral views.

At this later stage, the characteristic rope-ladder organization of the ventral nerve cord was disrupted in pax6KO larvae, with altered commissural organization and abnormal positioning of longitudinal connectives. These defects indicate that the early patterning alterations observed in *pax6* mutants are progressively translated into large-scale structural defects during nervous system maturation. Thus, *pax6* is required not only for the establishment of neural progenitor domains and neuronal marker expression, but also for the proper assembly of ventral nerve cord architecture.

### 3.6. A schematic model summarizes Pax6-dependent reorganization of mediolateral patterning

The changes observed in individual marker expression patterns were integrated into a schematic model of mediolateral ventral nerve cord organization (Fig. 6). In WT animals, *msx*, *pax3/7*, *pax6*, *nk6* and *nk2.2* occupy partially overlapping domains arranged in a stereotypical mediolateral order. Loss of *pax6* alters this organization by reducing the lateral *pax3/7* territory, narrowing the *nk6* domain and shifting the boundaries of the *nk2.2* domain. As a consequence, the relative proportions and overlaps of adjacent progenitor territories are modified.

**Fig. 6.**
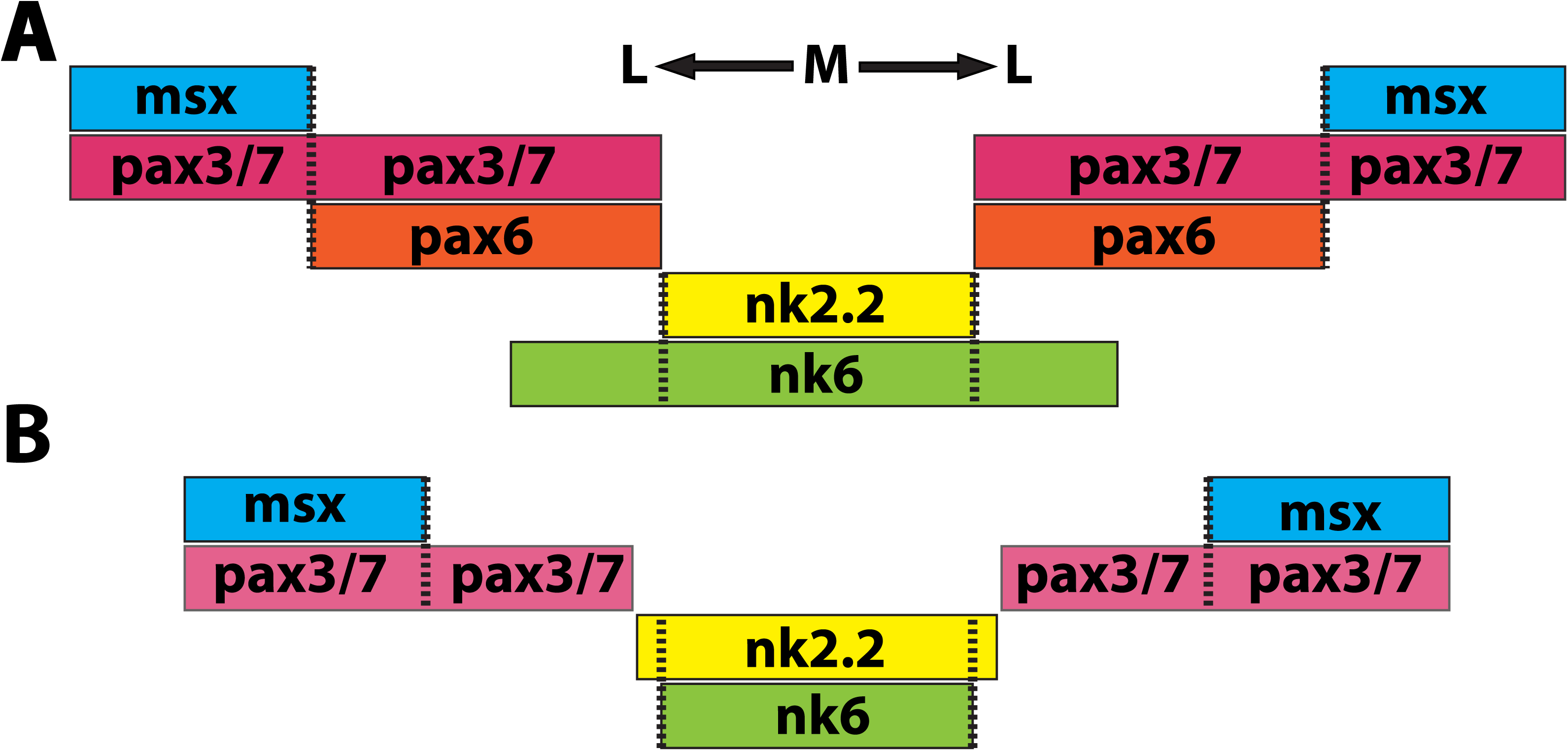
Schematic representation of mediolateral patterning of the ventral nerve cord in WT (A) and *pax6* KO (B) animals. Colored bars depict the relative expression domains of *msx, pax3/7, pax6, nk6*, and *nk2.2* and their overlaps along the mediolateral axis. Comparison of WT and KO animals illustrates the changes in domain organization observed in the mutant. Dashed lines indicate expression domain boundaries. M, medial; L, lateral.

This model emphasizes that the principal consequence of pax6 loss is not complete elimination of the dorsoventral patterning system, but reorganization of domain boundaries and domain overlaps. These boundary changes correlate with the later defects observed in serotonergic, cholinergic and motoneuron-associated markers, suggesting that Pax6 contributes to the correct balance between neighboring progenitor territories during ventral nerve cord development.

## 4. Discussion and Conclusions

This study identifies *pax6* as an important regulator of ventral nerve cord patterning and neuronal differentiation in the annelid *Platynereis dumerilii*. Using a ZFN-generated *pax6* mutant carrying a 61 bp deletion in the paired-domain coding region, we show that loss of pax6 alters the expression of several mediolateral patterning genes, including *nk2.2*, *nk6* and *pax3/7*, while *msx* remains largely unaffected. These early patterning abnormalities are followed by selective changes in neuronal markers at 48 hpf, broader defects in *TrpH*, *ChAT*, *VAChT* and *nk2.2* expression at 6 dpf, and disruption of the mature rope-ladder organization of the ventral nerve cord. Together, these findings support a model in which *pax6* acts high in the regulatory hierarchy that links mediolateral patterning of the ventral neuroectoderm to neuronal subtype specification and later nervous system morphogenesis.

*Pax6* is best known as a deeply conserved regulator of eye development, but it also has important roles in the developing nervous system across bilaterians (Callaerts et al., 2001; Halder et al., 1995; Hill et al., 1991; Kozmik and Kozmikova, 2024; Manuel et al., 2015; Osumi et al., 1997; Walther and Gruss, 1991). In vertebrates, *Pax6* is expressed in defined progenitor domains of the neural tube and contributes to the specification of ventral interneuron and motoneuron populations. It functions as part of a cross-regulatory network that includes Nkx2.2, Nkx6-class factors, *Dbx* genes, *Irx3* and *Olig2*, thereby converting graded morphogen inputs into discrete progenitor identities (Briscoe and Novitch, 2008; Briscoe et al., 2000; Ericson et al., 1997; Jessell, 2000; Kutejova et al., 2016; Vallstedt et al., 2001). The altered *nk2.2* and *nk6* domains observed in *Platynereis pax6* mutants are consistent with this general regulatory logic, even though the precise interactions in annelids remain to be tested directly.

In vertebrate neural tube patterning, domain boundaries are not simply passive readouts of morphogen gradients; they are actively sharpened by cross-repression between transcription factors expressed in neighboring progenitor domains (Balaskas et al., 2012; Briscoe et al., 2000; Dessaud et al., 2008; Vallstedt et al., 2001). A similar interpretation is supported by the *Platynereis* phenotype. In pax6KO animals, *nk2.2* and nk6 are not abolished, but their boundaries and relative positions are altered. This suggests that *pax6* contributes to the stabilization of neighboring gene-expression territories rather than acting only as a binary activator or repressor of individual markers. The reduction of *pax3/7* expression further indicates that *pax6* also influences more lateral territories, while the relative preservation of *msx* suggests that the most lateral domain is less dependent on *pax6* at the stages analyzed.

The temporal sequence of phenotypes is particularly informative. At 34 hpf, the primary defects involve progenitor-domain organization. At 48 hpf, selective abnormalities appear in *TrpH*-positive serotonergic neurons and *hb9*-positive motoneuron-associated domains. By 6 dpf, defects are evident in multiple neuronal markers and in the overall morphology of the ventral nerve cord. This progression is consistent with a causal relationship in which altered progenitor-domain boundaries lead to abnormal neuronal specification and, subsequently, impaired circuit architecture. Similar links between progenitor identity and neuronal subtype output are well established in the vertebrate spinal cord, where changes in *Nkx2.2*, *Nkx6.1*, *Pax6* or *Olig2* alter the production of V3 interneurons, motoneurons and other ventral neuronal classes (Briscoe et al., 2000; Briscoe et al., 1999; Ericson et al., 1997; Novitch et al., 2001; Sander et al., 2000).

The neuronal phenotypes observed in *Platynereis* also agree with previous work suggesting that annelid ventral nerve cord neurons arise from defined mediolateral territories. In *Platynereis*, staggered expression of *nk2.2*, *nk6*, *pax6*, *pax3/7* and *msx* has been proposed to subdivide the ventral neuroectoderm into domains comparable to those of the vertebrate neural tube and the *Drosophila* neuroectoderm (Denes et al., 2007). Single-cell and spatial analyses of the *Platynereis* larval nervous system further show that the 6 dpf ventral nerve cord contains molecularly distinct neuronal populations with combinatorial transcription factor profiles, including motoneuron-like populations associated with *nk6, hb9* and *islet* expression (Achim et al., 2015; Vergara et al., 2017). Our observation that *hb9*, *TrpH*, *ChAT* and *VAChT* expression are altered in *pax6* mutants supports the idea that mediolateral patterning domains are functionally connected to neuronal subtype organization in this annelid.

The disruption of acetylated tubulin organization at 6 dpf extends this conclusion from gene expression to nervous system morphology. The rope-ladder arrangement of the annelid ventral nerve cord depends on the proper positioning of longitudinal connectives and segmentally repeated commissures. In pax6KO larvae, the abnormal commissural organization and altered connectives indicate that patterning defects eventually affect the architecture of the nervous system as a whole. This phenotype is consistent with the broader role of *Pax6* in neural differentiation and axon organization reported in vertebrates and insects, where *Pax6* affects progenitor identity, neuronal differentiation, axon guidance and brain morphogenesis (Bel-Vialar et al., 2007; Furukubo-Ttokunaga et al., 2009; Huettl et al., 2016; Jones et al., 2002).

The schematic model in Fig. 6 summarizes the central interpretation of this study. In WT animals, the ventral neuroectoderm contains a stereotypical mediolateral arrangement of partially overlapping *msx*, *pax3/7, pax6*, *nk6* and *nk2.2* domains. Loss of *pax6* reorganizes this arrangement: the *nk6* territory narrows, *nk2.2* boundaries shift, and *pax3/7* expression is reduced. This model indicates that *pax6* is required for the relative positioning and balance of adjacent domains rather than for the formation of the entire ventral neuroectoderm. In this respect, the *Platynereis* phenotype resembles vertebrate neural tube mutants in which disruption of one transcription factor causes boundary shifts and changes in neuronal output without eliminating all dorsoventral patterning (Briscoe et al., 2000; Ericson et al., 1997; Vallstedt et al., 2001).

These findings are relevant to the long-standing debate on the evolutionary origin of centralized nervous systems. The staggered expression of *nk2.2*, *nk6*, *pax6*, *pax3/7* and *msx/msh* in vertebrates, *Drosophila* and *Platynereis* has been interpreted as evidence that a mediolaterally patterned nerve cord was present in the last common ancestor of bilaterians (Arendt et al., 2008; Arendt and Nübler-Jung, 1999; Denes et al., 2007). However, broader taxonomic sampling has challenged a simple homology model. In several bilaterian lineages, including xenacoelomorphs, rotifers, brachiopods and the annelid *Owenia fusiformis*, similar transcription factors do not always show the same relationship to neuronal subtype organization, leading to the proposal that bilaterian nerve cords may have evolved convergently in multiple lineages (Martín-Durán and Hejnol, 2021; Martín-Durán et al., 2018).

Our data do not resolve this debate on their own, but they add an important functional dimension. Previous comparisons often relied on expression patterns alone, which can be difficult to interpret because genes may be redeployed during evolution. Here, disruption of *pax6* causes coordinated changes in neighboring patterning domains, neuronal markers and ventral nerve cord morphology. Thus, at least in *Platynereis*, the conserved expression domains are not merely descriptive landmarks but components of an active developmental regulatory system (Tosches and Arendt, 2013). This strengthens the case that part of the gene regulatory logic underlying mediolateral nerve cord patterning is deeply conserved, even if the extent of conservation varies among bilaterian lineages.

Functional data from other lophotrochozoans are still limited but broadly consistent with an important neural role for *pax6*. In the annelid *Capitella teleta*, pax6 knockdown disrupts eye and nervous system development, including defects in neuronal number, axon pathfinding and nerve positioning (Klann and Seaver, 2019). In planarians, *pax6*-related genes contribute to eye regeneration and neural patterning, while staggered expression of *nk2.2*, *nk6* and *pax6* has been described in the nervous system of *Schmidtea mediterranea* (Pineda et al., 2002; Wang et al., 2016). In the cephalopod *Sepia officinalis*, *nk2.1/nkx2*, *pax6*, *pax3/7* and *msx* expression domains have also been interpreted as evidence for conserved dorsoventral neural patterning within Spiralia (Buresi et al., 2016). Together with our mutant analysis in *Platynereis*, these data suggest that *pax6*-dependent neural patterning may be a recurrent and functionally important feature across several spiralian lineages.

At the same time, caution is warranted. The relationship between conserved transcription factor expression and nervous system homology is complex. Hemichordates provide a striking example: direct-developing *Saccoglossus* has a broadly distributed ectodermal nervous system and uses BMP signaling differently from vertebrates, whereas indirect-developing hemichordates show stronger evidence for BMP-dependent dorsoventral neural patterning (Lowe et al., 2006, 2015; Lowe et al., 2003; Su et al., 2019). Such comparisons indicate that ancestral developmental programs can be modified extensively, redeployed, or partially lost. Therefore, functional tests in additional lineages will be necessary to determine which aspects of the dorsoventral patterning network are ancestral for Bilateria and which evolved independently or were secondarily modified.

In conclusion, our results show that *pax6* is required for proper mediolateral patterning, neuronal marker expression and ventral nerve cord morphology in *Platynereis dumerilii*. The *pax6* mutant phenotype provides functional support for the idea that the annelid ventral nerve cord is patterned by a conserved regulatory system involving *pax6, nk2.2, nk6, pax3/7* and *msx*. By linking early progenitor-domain organization to later neuronal subtype and morphological defects, this study strengthens the view that conserved dorsoventral patterning factors have an active developmental role in annelid nervous system formation. More broadly, these findings contribute functional evidence to discussions of bilaterian nervous system evolution and highlight *Platynereis* as a valuable model for testing the conservation and diversification of neural patterning mechanisms.

## CRediT authorship contribution statement

**Jovana Doderovic:** Writing – original draft, Methodology, Investigation, Data curation.

**Miroslava Kolkova:** Methodology, Investigation, Data curation

**Anna Zitova:** Methodology

**Iryna Kozmikova:** Writing – review & editing, Data curation. Conceptualization.

**Zbynek Kozmik:** Writing – review & editing, Re- sources, Project administration, Methodology, Investigation, Funding acquisition, Data curation, Conceptualization.

## Acknowledgements

This work was supported by the institutional funding (RVO68378050-KAV-NPUI), grant by the Czech Science Foundation GA24-12482S (to Z.K.). We acknowledge the Light Microscopy Core Facility, IMG, Prague, Czech Republic, supported by MEYS – LM2023050, MEYS – CZ.02.1.01/0.0/0.0/18_046/0016045 and MEYS – CZ.02.01.01/00/23_015/0008205, for their support with the confocal imaging presented herein. We thank Veronika Noskova for technical assistance and Gaspar Jekely for plasmids used for WMISH probe generation.

